# Contribution of structural variation to genome structure: TAD fusion discovery and ranking

**DOI:** 10.1101/279356

**Authors:** Linh Huynh, Fereydoun Hormozdiari

**Affiliations:** Genome Center, UC Davis; UC Davis MIND institute; Department of Biochemistry and Molecular Medicine, UC Davis

## Abstract

The significant contribution of structural variants to function, disease, and evolution is widely reported. However, in many cases, the mechanism by which these variants contribute to the phenotype is not well understood. Recent studies reported structural variants that disrupted the three-dimensional genome structure by fusing two topologically associating domains (TADs), such that enhancers from one TAD interacted with genes from the other TAD, and could cause severe developmental disorders. However, no computational method exists for directly scoring and ranking structural variations based on their effect on the three-dimensional structure such as the TAD disruption to guide further studies of their biological function. In this paper, we formally define TAD fusion and provide a combinatorial approach for assigning a score to quantify the level of TAD fusion for each deletion denoted as TAD fusion score. We also show that our method outperforms the approaches which use predicted TADs and overlay the deletion on them to predict TAD fusion. Furthermore, we show that deletions that cause TAD fusion are rare and under negative selection in general population. Finally, we show that our method correctly gives higher scores to deletions reported to cause various disorders (developmental disorder and cancer) in comparison to the deletions reported in the 1000 genomes project.

## 1 INTRODUCTION

The genome and chromosomes of all species encode the genetic information for the production of RNA and proteins and it is represented as a linear sequence. Genes and other regulatory elements are organized and positioned in this linear space. However, in reality, the chromosomes in each cell are folded as a complex three-dimensional (3D) structure. The “interactions” or physical connection between different segments of the genome in this 3D structure regulates the whole transcriptional process of all genes [1, 2]. The 3D organization and folding of chromosomes govern how the distal genomic elements interact by bringing them into direct contact with each other. These distal functional elements such as enhancers are the main source of controlling the gene expression regulation in various tissues, and their interactions are the governing force of these regulations. The folding and interaction between genomic elements also control cellular processes such as transcription, replication and DNA damage repair [3, 4].

The first in-depth studies on the 3D structure of chromosomes were obtained using microscopy techniques [5]. However, these approaches tend not to be able to resolve the physical interactions and the 3D structure in highresolution and high-throughput fashion [6, 1]. With the advent of various biomolecular chromosome conformation capture techniques in the past few years, our understanding of the 3D structure of genomes and its contribution to various biological properties have been revolutionized [6, 1, 7, 8]. These chromosome conformation capture techniques are now capable of capturing chromosomal contacts in different conditions, diverse throughput and resolutions. Some of the most popular techniques developed including chromosome conformation capture (3C), circular chromosome conformation capture (4C) and chromosome conformation capture carbon copy (5C) [9, 10, 11, 12].

Recently, an extension of the 3C technique has been developed which is capable of producing genome-wide interaction of all chromosomes in high-throughput and high-resolution fashion denoted as Hi-C [13, 14, 15, 16]. Hi-C technique that is based on proximity ligation and pair-end sequencing allows researchers to capture the 3D structure of the chromosomes at a kilobase pair (kbp) scale. This technique produces a genome-wide sequencing library that provides a proxy for measuring the three-dimensional distances among all possible locus pairs in the genome [17]. Provided the Hi-C experiment paired-end reads the next phase is producing the (normalized) contact frequency maps/matrix. This contact matrix provides a qualitative measurement of physical closeness and interaction of different segments of the genome in 3D structure.

Hi-C is a powerful technique which can provide the researchers with valuable biological insights such as the 3D structure of genome, genomic physical interactions and chromatin contacts. One of the most interesting novel discoveries using Hi-C data is the compartments of the genome into hundreds of kilobase pair segments which interactions are highly enriched inside each segment and are significantly depleted between adjacent segments. These segments are denoted as topological associated domains (TADs) and can be seen as continuous square domains on the diagonal of the Hi-C contact matrix [4]. It is shown that TAD boundaries are significantly enriched with insulator proteins. These proteins are shown to be able to block interactions between different regulatory elements and are the main reason there is significant depletion of interactions between two adjacent TADs. The dominant insulator protein that acts as TAD boundaries in mammalian cells is CTCF [4, 3]. TADs are hypothesized to be conserved between different cell types and across close species, however it is not trivial to quantify this mainly due to difficulties of accurate TAD discovery, existence of sub-TADs and nested TADs [18, 19, 20].

In a recent experiment, Dali and Blanchette[20] manually annotated TAD structures in a randomly selected segments of genome and compared against TAD prediction using various computational tools. They reported a significant discordance between various prediction algorithms. Furthermore, it was observed that tools generally had low sensitivity, often picking up less than 10% of manually annotated TAD structures. In fact, almost 25% of the manually annotated boundaries never got detected by any of the tools [20]. In another study [18] similar disagreement between computational tools for TAD prediction was observed. Furthermore, TAD callers returned different numbers of TADs with different mean sizes [18]. Existence of sub-TADs, nested-TADs [19, 20], difference of TAD structure for cells in the same tissue and limiting assumptions made by different computational methods are some of the main reasons for these discrepancies.

In a seminal paper by Lupianez et al. [21] it was shown that structural variation which disrupted TADs (by deleting TAD boundaries) can result in novel enhancer-promoter interactions which in return can cause severe developmental disorders. More specially, it was shown that structural variations which disrupted the TAD boundary (i.e., the CTCF insulators) of WNT6/IHH/EPHA4/PAX3 locus fused two TADs which caused malformation syndromes [21]. Similarly findings have shown that structural variations (SVs) that cause TAD organization disruption and fusion of TADs in different loci can cause various developmental disorders [21, 22, 23, 24, 25, 26]. In addition to the developmental disorder, recent findings reported similar TAD disruptions in cancer cells [27, 28, 29] and an effective mechanism for activation of oncogenes. It was shown that TAD fusion could result in activation of oncogenes by creating novel genomic interactions between enhancers and an oncogene. For example, it was shown that deletion of TAD boundaries resulted in activation of oncogenes *TAL1* or *PDGFRA* which contributed to cancer [27, 29, 28, 30].

These studies have shown the importance of being able to study the impact of structural variations on the threedimensional genome organization. Specifically, if a deletion results in the disruption of two or more TADs by fusing them and creating novel genomic interactions (figure 1). However, no method exists for directly scoring and ranking deletions based on their effect on 3D structure and TAD disruption. In this paper, we formally define TAD fusion and provide a combinatorial approach for assigning a score to quantify the level of TAD fusion for *each deletion* denoted as “TAD fusion score”. We also show that in practice our approach is superior for predicting deletions which result in TAD fusion in comparison to approaches which directly use predicted TADs. Furthermore, we show that deletions which cause TAD fusion are rare and are negatively selected against in general population. Finally, we show that our method correctly gives significant higher scores to the deletions reported to cause various disorders (developmental disorder and cancer) as a result of disrupting 3D structure in comparison to the deletions reported in the 1000 genomes project.

**Figure 1:**
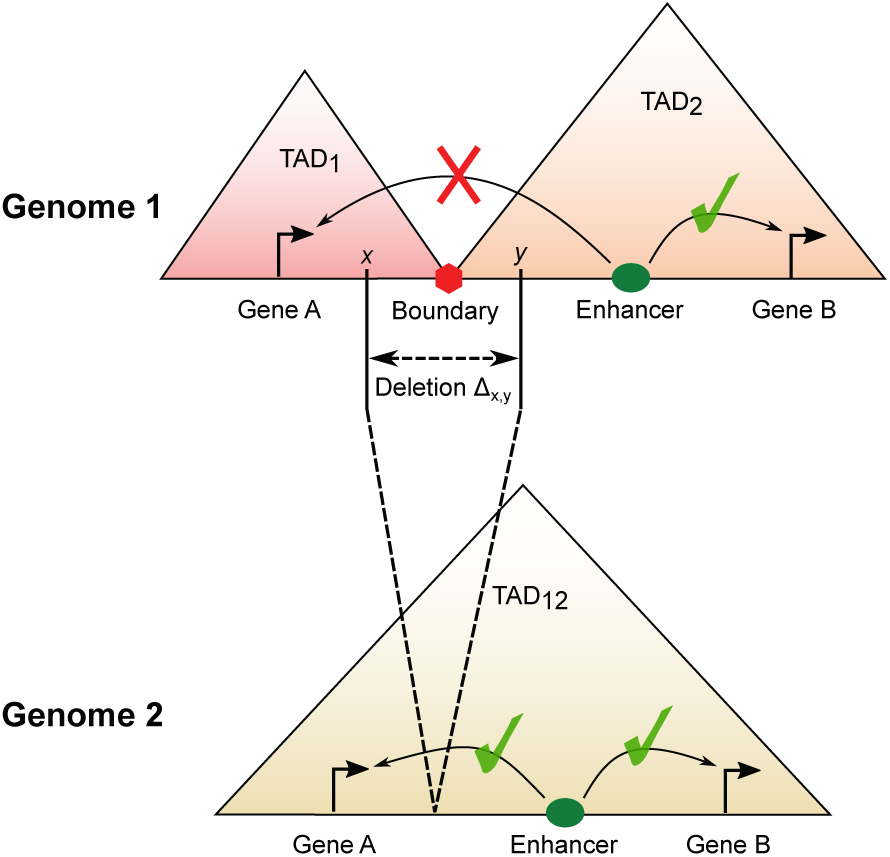
An illustration of the TAD fusion where a structural variation can affect the genome structure.

## 2 RESULTS

### 2.1 Methods Overview

To study TAD fusion and its contribution to disease and evolution we will define a novel score to assign to any structural variation (SV) based on its level of modifying the 3D genome structure and potential of causing a TAD fusion. We also develop a novel computational method to calculate this TAD fusion score (the details can be seen in section 5) for any deletions. In this paper, we are mainly considering the **deletion** as the input structure variation, but similar idea can be extended to consider inversions or translocations.

**TAD fusion score problem and framework:** Given a Hi-C contact frequency matrix of a chromosome and the coordinates of a deletion, we define a TAD fusion score as *the expected total number of changes in pairwise genomic interactions as a result of the deletion*. The input to the method consists of the Hi-C matrix of the genome with reference allele (i.e., without the deletion) and the coordinates of the deletion of interest in the test genome. The main output of the method is a score representing the expected number of changes in the pairwise genomic interactions as a result of the deletion. We denoted this as the **TAD fusion score** of the deletion.

We are proposing a **two-step framework** for calculating the TAD fusion score: (i) predicting the new Hi-C contact matrix *G* of the mutated genome (i.e. with the deletion), given as input the Hi-C contact matrix *H* of a genome without the deletion and the deletion coordinates; (ii) comparing the predicted Hi-C contact matrix *G* with the original Hi-C contact matrix *H* to estimate the changes in number of pairwise genomic interactions as a result of that deletion.

### 2.2 Data

We used Hi-C data of the human cell GM12878 from *in situ* experiments [31]. We chose the resolution 5kbp since the TAD structure conservation (with other cell lines) was validated at that resolution. For deletions, we used the reported deletions in the 1000 genomes project (1KG) [32], the deletions reported in tumor samples from The Cancer Genome Atlas (TCGA https://cancergenome.nih.gov/) project and a small set of deletions reported and validated to cause developmental disease by disrupting genome structure [21, 24, 22, 23]. Although no method existed for explicit prediction and scoring of SVs for their disruption of genome structure and causing TAD fusion for comparison, we considered the boundaries predicted by several state of the art TAD callers [33, 34, 35, 31] for TAD fusion prediction. The setting of these methods is presented in table S1. For evaluation we also considers the overlap of the predicted TAD fusions with reported CTCF binding sites in GM12878 samples from ENCODE data. All data (Hi-C, 1KG deletions, deletion mutations of GM12878, CTCF binding sites) were aligned with the reference human genome b37 (hg19). As such, 82 deletion mutations of GM12878 were removed from 1KG data from consideration as they were also reported in the sample GM12878. Since we used Hi-C data at the resolution 5kbp, only deletion mutations that were longer than 10kbp were considered to ensure each deletion removed completely at least one 5kbp-bin. There were a total of 7383 such deletion reported in the 1000 genomes project [32].

**Evaluating the Hi-C contact frequency prediction (Step 1)**: At the core of our proposed method to estimate the TAD fusion score is to predict the Hi-C contact frequencies as a result of the deletion (i.e., the step 1 of the method as detailed in method sections 5.1). We have denoted our proposed approach for predicting the new contact frequencies as the **distance based model** (See section 5.1). We have shown our *distance based model* for predicting Hi-C contact frequencies outperforms the traditional model based on the genomic length [16] for Hi-C contact frequency predictions using both GM12878 data (see Supplementary Material Section 1.1, Supplementary Figure S1) and cancer cell line K562 (see Supplementary Figure S2 and Supplementary Table S2).

### 2.3 Evaluating the TAD fusion score

The main objective of the approach presented here is to assign a score to each SV based on the potential to disrupt the genome structure and causing TAD fusion. We compared our TAD fusion scoring approach with four state of the art methods for TAD prediction and used their boundary prediction to predict TAD fusion. For TAD callers we assumed that a deletion is causing TAD fusion based on their TAD calls if the two breakpoints of the deletion are inside two different TADs (i.e., the TAD caller is predicting that specific deletion is deleting a TAD boundary).

Note that, there does not exist a truth set to compare TAD fusion calls made using various TAD callers to our novel fusion scores calculated. Thus, we used the consensus of the TAD caller predictions as the indication of ground-truth. We compared our TAD fusion scores against the TAD fusions predicted by each of the four methods separately by using the majority of the fusions predicted by the three remaining methods as the truth set. Note that, for each TAD caller that predicted a set of deletions as TAD-fusion in the 1000 genomes deletion set, we considered the same number of deletions with highest TAD fusion score from our proposed approach for comparison. We repeated this approach for each of the four TAD callers we considered for comparison. Figure 2.a shows that there is higher agreement between our method with the consensus prediction than any other tool.

**Figure 2:**
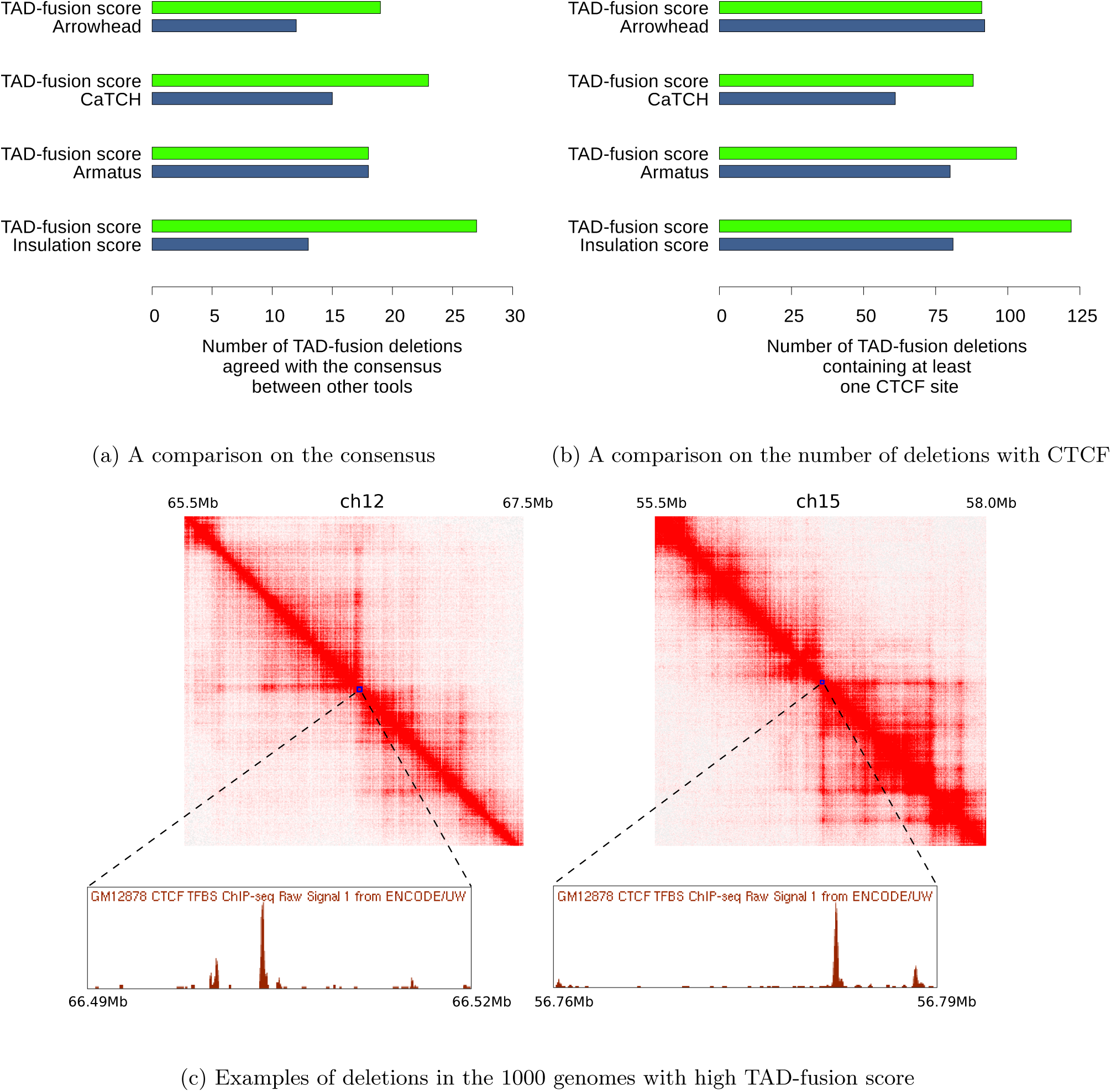
A benchmark on the TAD-fusion prediction with deletions from 1000 genomes project. To compare our method with a method that outputs *k* predicted TAD-fusion deletions, we assign deletions that the TAD-fusion score is in top *k* as TAD-fusion deletions. (a) A comparison on the agreement with the consensus (i.e. a deletion is a TAD-fusion one if there are at least two other tools predict that), (b) a comparison on the number of TAD-fusion deletions that contain at least one CTCF binding site, (c) Examples of deletions (from 1000 genomes project) that have a very high TAD-fusion score but are not predicted as TAD-fusions by any other tools predicting TADs. Note that both these deletions clearly are deleting CTCF binding sites (i.e. potential TAD boundaries).

A stronger evidence of advantage of our TAD fusion score prediction in comparison to using available TAD callers for predicting fusion comes from the CTCF binding sites in the genome. There are clear evidence that CTCF binding sites play a major role in the genome organization and they are very significantly enriched at that TAD boundaries [3, 4]. Hence, the deletion of these binding sites has a higher chance of leading to the TAD structure disruption and TAD fusion [21, 36, 37, 38]. Thus, we evaluate the number of predicted TAD fusions that contain at least a CTCF binding site (figure 2.b). The top set of deletions selected by our TAD fusion score approach outperformed approaches which were based on TAD callers. Note that, we did not take into account any information about the CTCF sites in the model or the prediction.

Finally, we also (visually) investigated few of the 1000 genomes deletions with very high TAD-fusion score which were not predicted by any of the TAD callers to have a boundary inside these deletions. Figure 2.c illustrates two such examples that are rare and relatively small deletions from the 1000 genomes project. These two deletions were assigned a very high TAD fusion score by our method (in the top 100 deletions), however were not predicted to have a TAD boundary by any other tool (CaTCH, Amaratus, Arrowhead and InsulationScore) we tested. We specifically selected these two deletions to depict here as they tend to be in the lower range of the length of the deletion (26kbp and 32kbp). In both deletions based on the visual inspection of the Hi-C data and the CTCF binding sites reported, it seems to support our TAD fusion scoring.

### 2.4 TAD fusion is under negative selection in healthy samples

In few recent studies, it was shown that SVs which disrupted genome structure and caused TAD fusion could result in various developmental disorders. Thus, it is expected that TAD fusion events should be negatively selected in normal population. For testing this hypothesis we compare the TAD fusion scores calculated for deletions reported in the 1000 genomes project against a null model where the same set of deletions are randomly assigned in genome. We randomized the permutation of the 1000 genomes project deletions to ensure the same number and length of deletions per chromosome and calculated the TAD-fusion score per permuted deletion and repeated this permutation procedure for 5, 000 times. We then compared deletions in the 1000 genomes project with TAD fusion score above various cut-offs (table S3) and observed the number of such deletions was significantly less in the 1000 genomes project set in comparison with the set of permuted data (if we set the cut-off at the top 100, we got a p-value *<* 10^−37^, figure 3.a). We then compared the sum of TAD fusion scores calculated for deletions in the 1000 genomes project against the permuted set and observed that the 1000 genomes deletion set had significantly smaller fusion score sum than the random deletions had (p-value *<* 10^−19^, figure 3.b). These results indicate that deletions which potentially cause TAD fusion are significantly less in the normal genome population. We further investigated the correlation between the allele count of reported deletions in the 1000 genomes project against the calculated TAD fusion score. We observed that deletions with higher TAD fusion score have significantly lower allele count. To better visualize we grouped the 1000 genomes project deletions into bins based on the calculated TAD fusion score and compared against the allele count (figure 3.c) with Spearman rank correlation of *r <* − 0.83 and *p <* 0.00064. Note that deletions with highest TAD fusion score (*>* 500) the average allele count in the 1000 genomes samples is almost 1 (the lowest possible value). Thus, indicating that deletion with high TAD fusion score are under negative selection.

**Figure 3:**
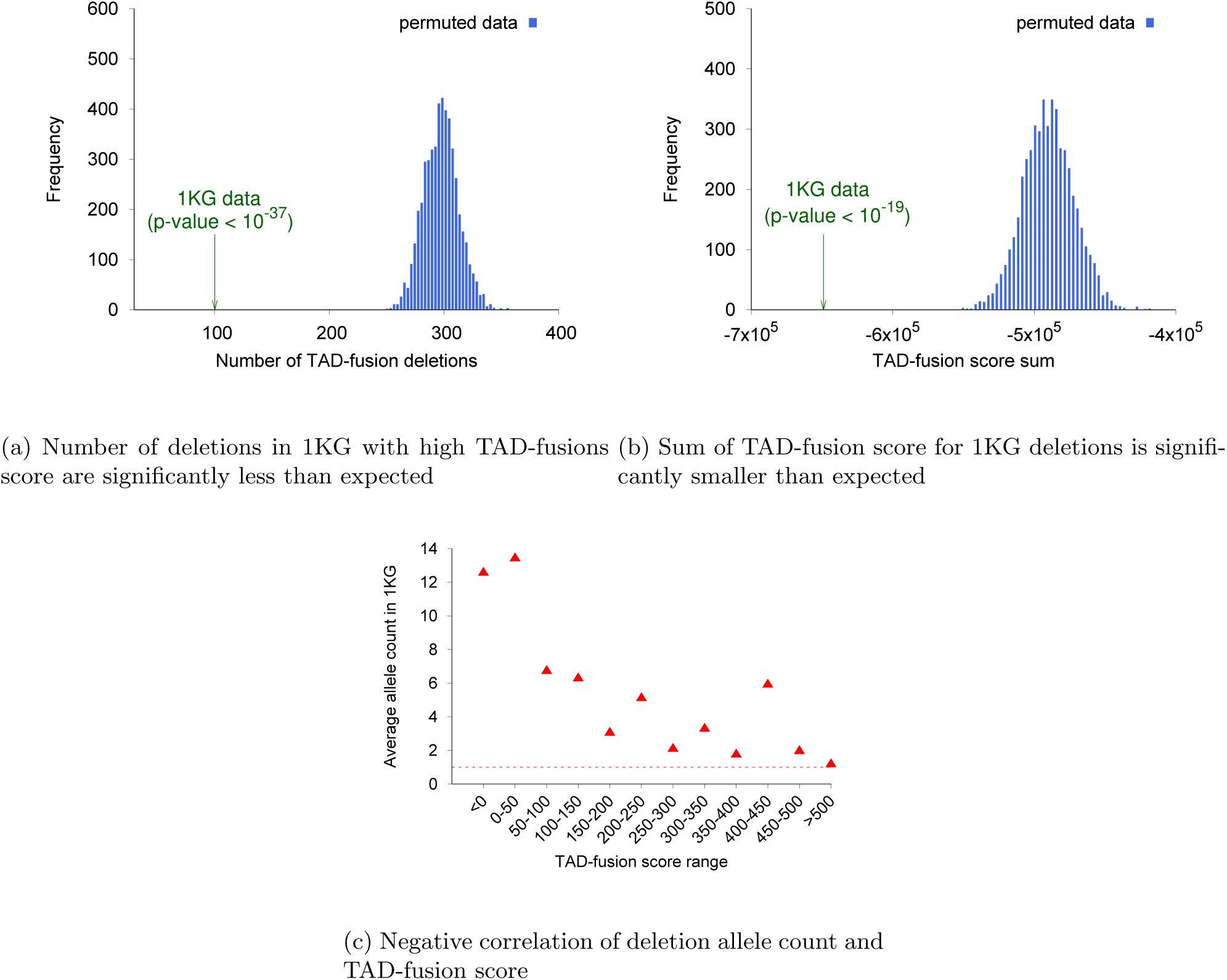
TAD-fusion score of the deletion in the 1000 genomes project (1KG). (a) The number of TAD-fusion deletions (top 100) of 1KG data and permuted data (5000 random samples), (b) The TAD-fusion score sum of all deletions of 1KG data and permuted data (5000 random samples), (c) The average allele count of 1KG deletions (deletions are binned by the TAD-fusion score).

### 2.5 Contribution of TAD fusion to diseases

For studying the contribution of TAD fusion to human disease, We utilized a list of nine deletions which were validated to cause various disorders as a result of genome structure disruption [23, 21, 22, 24]. We estimated the TAD fusion score for these nine deletions with comparison to the deletions reported in the 1000 genomes project. The result shows that the TAD fusion score of these nine deletions is significantly higher than almost all reported deletions in the 1000 genomes project (figure 4.a). This supports the claim that our method assigns a higher scores to deletions which disrupt genome structure and cause TAD fusions. We further investigate these nine deletions by partitioning them into a series of smaller 20 kbp deletions (with 10 kbp overlap). We then calculate the TAD-fusion score for all these 20 kbp deletions covering the original deletions. We first depict the predicted TAD-fusion score for these 20 kbp deletions produced from two of the disease deletions with the matching Hi-C maps (Figure 4.b note that very similar results was obtained for other deletions tested). In addition to the TAD-fusion score for each of these 20 kbp deletions the CTCF binding sites inside each of these deletions was also depicted (Figure 4.b). We can clearly see in Figure 4.b that the 20 kbp deletions which span the boundaries of the TADs have gotten a much higher TAD-fusion score, which also match with the known CTCF binding sites (Figure 4.b). We further investigate the TAD-fusion score for these 20 kbp deletions which include a known CTCF binding site for all the nine disease deletion. We observed that a statistically higher TAD-fusion score is given for the 20 kbp deletions which remove a CTCF binding site versus the ones that do not remove CTCF binding site (Figure 4.c).

**Figure 4:**
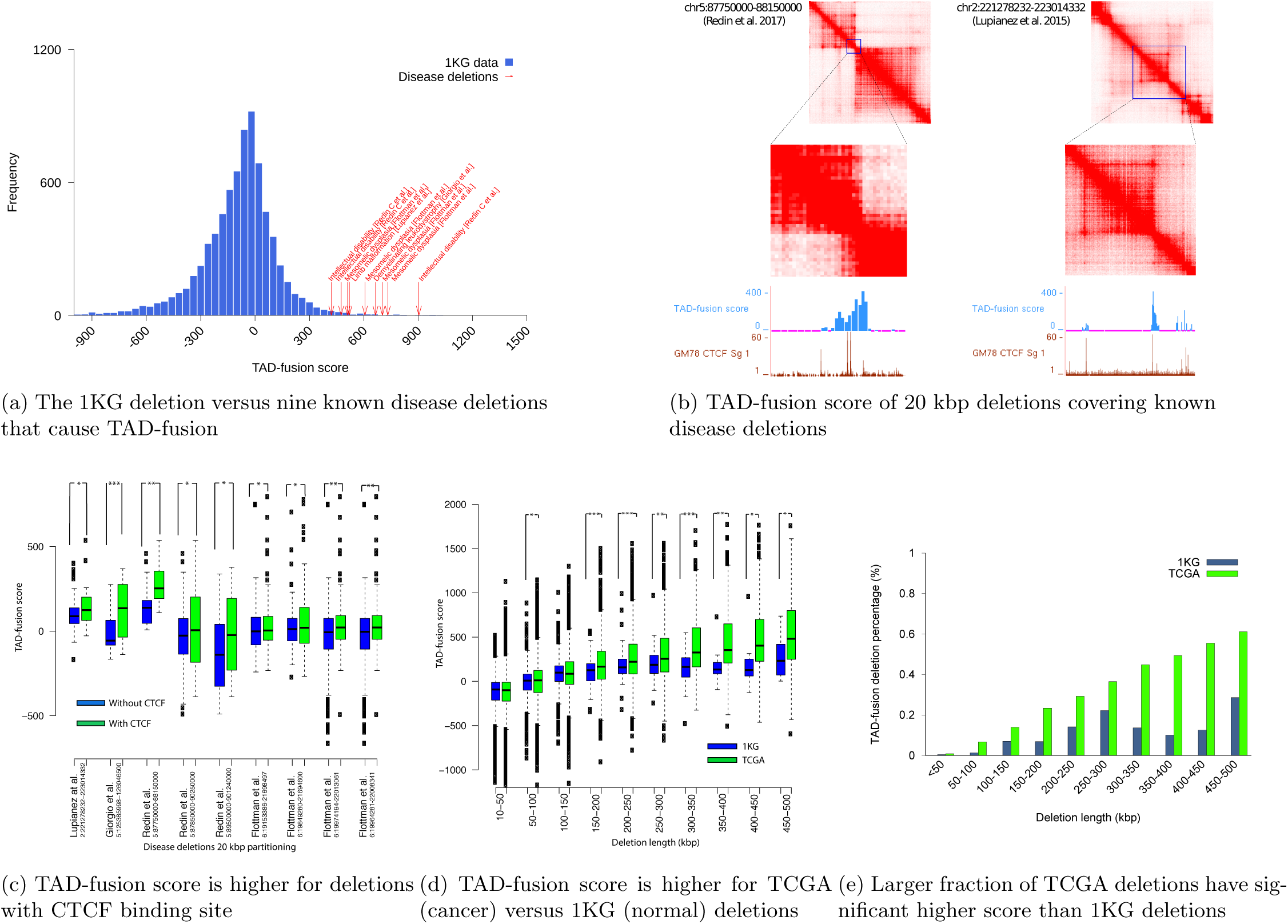
(a) The TAD-fusion score of 1KG deletions and nine reported disease deletions that cause disorders by causing TAD disruption (i.e. TAD-fusion). (b) TAD-fusion score and CTCF binding sites for 20 kbp deletions spanning two of the known disease deletion and the respective Hi-C contact maps. Clear correlation for the TAD-fusion score for deletions which span the TAD boundaries and have CTCF binding sites can be seen. (c) Significant higher TADfusion score for 20 kbp deletions which remove a CTCF binding site versus the ones that do not. (d) Comparison of TAD-fusion score between cancer deletions (TCGA) versus deletions in normal samples (1KG) (e) A comparison between fraction of deletions in 1KG deletion set and TCGA deletion set which have a high TAD-fusion score.

We also explored the TAD-fusion score for deletions reported in tumor cells from TCGA (The Caner Genome Atlas) and compared them against deletions from the 1000 genomes project of similar length. We observed that almost for all different length of deletions the TCGA deletions have a significantly higher TAD-fusion score than the 1000 genomes deletions (Figure 4.d and Figure 4.e).

## 3 Discussion

Chromosome folding and structure in 3-dimensional space plays an important role in gene expression and regulation. Advance molecular biology techniques such as the Hi-C protocol provide experimental evidence that the chromosome is organized in modular hierarchical domains such as compartments, TADs and sub-TADs. Motivated by few case studies of limb development and developmental disability, it is hypothesized that deletions (and other SVs) that lead to 3D structure disruption and contribute to the etiology of these disorders. In most cases, these deletions contribute to the disorder by fusing two TADs. Thus, these types of deletions should be negatively selected for and they should be rare in normal samples. Although, there are many studies on Hi-C data analysis such as normalizing the raw data, finding significant interactions or identifying the TADs, there is no method that can directly predict if a SV results in a TAD fusion. In this study, we have developed a novel computational method for assigning TAD fusion score to any input deletion based on its potential contribution to fusing TADs. As part of this method, we solve two related subproblems: (i) The first problem is how we can predict the changes to Hi-C data due to a deletion in the genome, (ii) and the second problem is how we can compare two Hi-C contact matrices to find the significant difference between them. For the first problem of predicting the changes in Hi-C data due to a deletion, more accurate model is the key to improve the prediction. The distance based model presented here agrees well with the experimental data by taking into account 3D distance features. However, it can still be improved by considering the mechanism of creating interactions, such as the loop extrusion model [39, 40] or the slip link model [41]. Furthermore, the model could also take into account other biological features such as histone modification marks, DNA methylation or DNA accessibility [34, 42]. In our prediction algorithm, we assume that the deletion only changes the interactions that cross the deletion (i.e. between two bins, one is in the upstream and one is in the downstream of the deletion). However, this constraint may not always hold and the model should be extended to consider also the changes between bins at one side of the deletion. For the second problem, we can evaluate the change of only enhancer-promoter interactions rather than the interaction between all bins. In addition, the comparison on the contact frequency should also be normalized for each specific pair, gene, or region to achieve better result. Other cell lines also need to be evaluated although the TAD structure may be conserved. Further experiments are needed to validate our predicted results. For example, we can make a deletion in the genome (e.g. with CRISPR) and generate the new Hi-C data and also measure the gene expression change. These gene expression may not change constantly since the regulation depends not only on the genomic interactions but also on other factors (e.g. presence or absence of transcription factors), thus we need to capture the gene expression dynamically for the validation. However, this obstacle could be overcome since more data are contributed by the community. In addition, due to the complexity of the original optimization problem, we have applied the approximation for some steps, any improvement on this approximation to make it closer to the original one may improve the result. Since the LP solver still has a large complexity, improvements or adding more constraint is necessary. The number of variables can be reduced by simplifying the objective function such as removing less significant location pair while adding larger weight for important ones. Other metric rather than L1 may also need to be evaluated. Further, the model is fitted in the log scale and thus zero entries should be normalized, this problem is similar to the problem of normalizing drop-out data in processing single cell gene expression data. There are several applications of the proposed method for TAD-fusion discovery, it will provide biologists with a way to rank and pick deletions which are potentially causing significant disruption for genome structure. Furthermore, TAD-fusion discovery will provide a novel mechanistic explanation of how a group of non-coding deletions (and SVs in general) are contributing to developmental disorders or cancer Finally, the approach presented here for TAD fusion scoring of deletions can be extended to also consider other types of structural variants, such as inversions and translocations.

## STAR METHODS

### 4 KEY RESOURCES TABLE

| Resource | Source | Identifier |
| --- | --- | --- |
| Software |  |  |
| Genome browser |  | https://genome.ucsc.edu |
| ILOG CPLEX |  | http://www.ilog.com/products/cplex |
| Insulation-score | [35] | https://github.com/dekkerlab/crane-nature-2015 |
| CaTCH | [34] | https://github.com/zhanyinx/CaTCH_R |
| Armatus | [33] | https://github.com/kingsfordgroup/armatus |
| Arrowhead | [31] | GSE63525 |
| TAD-fusion score | This paper | https://github.com/huynhvietlinh/FusionScore |
| Other |  |  |
| HiC data | [31] | GSE63525 |
| HiCCUP loops | [31] | GSE63525 |
| K562 deletion data |  | https://www.biorxiv.org/content/early/2017/12/02/192344 |
| CTCF binding sites | [43] | https://www.encodeproject.org |
| 1KG deletion data | [32] | http://www.internationalgenome.org/data |
| TCGA deletion data | TCGA | https://portal.gdc.cancer.gov |

### 5 METHODS DETAILS

Our goal is to develop a *computational method to provide a score to any structural variation (SV) based on its level of modifying the 3D genomic structure and potential of causing a TAD fusion.* We are assuming the input to the method consists of the Hi-C matrix of the genome with reference allele (i.e., without the SV) and the coordinates of the SV, and the output is a score representing **the number of new genomic interactions** made (i.e., TAD fusion score) as a result of the SV. For this paper, SVs are assuming only homozygous and non-overlapping deletions however this approach can be extended to consider other SV types (e.g. translocations, inversions).

**Definitions and notations:**

- Chromosome location bin: Considering the limited sequencing coverage that produces the Hi-C data, achieving single base-pair resolution is impossible. Thus, it is common in practice to partition the genome into nonoverlapping segments (bins) to summarize the data. The contact frequency map is the summarization of each chromosome by partitioning this chromosome into non-overlapping bins of the same length (e.g. 5kb), and calculate the average number of contacts between any two bins. We assume that we have *n* bins which are numbered across the chromosome as 1, 2, 3, *…, n*.
- Genomic interaction: Two bins *i* and *j* are considered interacting if they create a physical interaction in 3dimensional space. Some of the well-known examples of such interactions are the enhancer-promoter interactions. Our goal is to calculate the *expected number of new interactions* created as a result of a SV (e.g., deletions).
- Hi-C contact frequency map/matrix: Let *H ∈ R*^*n×n*^ be a symmetric matrix constructed from Hi-C data where *H*_*i,j*_ is the average contact frequency between *i*^*th*^ bin and *j*^*th*^ bin (i.e. *H*_*i,j*_ is large if *i* and *j* are close in 3-dimensional space, otherwise, *H*_*i,j*_ is small if they are not close in 3-dimensional space). Note that, if two bins *i* and *j* are interacting with each other they have to be very close in 3-dimensional space.
- Deletion structural variation: We denote Δ_*x,y*_ as the deletion from the *x*^*th*^ bin to the *y*^*th*^ bin of a chromosome. We round the coordinates down to calculate the location bin of a deletion.

**TAD fusion prediction problem:** Given a Hi-C contact frequency matrix of a chromosome and the coordinates of a deletion SV, we predict if this SV results in a TAD fusion and assign a TAD fusion score to that SV. We define a TAD fusion based on new genomic interactions created between different bins as a result of the SV. By that, **TAD fusion score** is defined as the *expected number of additional genomic interactions created* as a result of the SV.

We are proposing a two-step framework for calculating the TAD fusion score of a homozygous deletion: (i) predicting a new Hi-C contact matrix *G* of the mutated chromosome (i.e. with the deletion) given the Hi-C contact matrix *H* of a genome without the deletion and the deletion coordinates as the inputs; (ii) comparing this predicted/new Hi-C contact matrix *G* with the original Hi-C contact matrix *H* to estimate the number of new interactions created as a result of that deletion SV. In the next two subsections, we introduce the algorithms we have developed as part of this framework to solve each of these two subproblems.

#### 5.1 Predicting contact frequency matrix resulting from a deletion

In this subsection, we provide a combinatorial algorithm for predicting the contact frequency matrix *G* of a genome with a homozygous deletion Δ_*x,y*_ given the Hi-C contact frequency matrix *H* of the genome without the deletions as input. We assume that the contact frequency between any two bins *i, j* will not be changed as a result of the deletion if both bins *i* and *j* are in the upstream or the downstream of the deletion (i.e. *G*_*i,j*_ = *H*_*i,j*_ if 1 *≤ i, j < x* or *y < i, j ≤ n*). Our goal is to predict the contact frequency *G*_*i,j*_ between any bin *i* in the upstream of the deletion and any bin *j* in the downstream of the deletion (i.e., 1 *≤ i < x* and *y < j ≤ n*).

##### 5.1.1 Modeling the contact frequency

In this subsection, we introduce the main models for predicting Hi-C contact frequencies between any pair of bins based on the genomic or physical distance between the bins [44, 45, 46, 47]. It has been shown that power-law scaling based on the genomic distance or physical distance best captures the contact frequencies observed in Hi-C experiments [44, 45, 46]. We denote the power-law scaling factor by parameter *β* and model the Hi-C contact map as 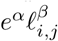 [44, 46]. Recent studies have shown that the power-law scaling *β* that best matches the Hi-C contact frequencies is approximately equal to −1 [48]. We denote this simple model of Hi-C contact frequencies as the *genomic length based model*. Furthermore, it is shown that contact frequency between two bins is also effected by the genomic properties of these two bins (e.g. GC content, mappability, etc). Therefore, in predicting the Hi-C contact frequencies using the genomic distance, we also introduce a parameter *α*_*i*_ for each bin *i* to capture the genomic properties/biases of this bin. Thus, the genomic *length based model* to predict the contact frequency for Hi-C data can be extended to 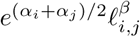 [47, 46].

Although the genomic length based model for predicting Hi-C contact frequencies is very useful, it is shown recently that a *metric distance* which captures the physical distance would be a better predictor of the Hi-C contact frequencies observed [47]. We denote this metric distance which represents the physical distance between bins *i* and *j* as *d*_*i,j*_. Similarly it is shown that the Hi-C contact frequencies obey a power-law scaling of the physical distance and is dependent on the genomic properties at bins *i* and *j* (i.e. *α*_*i*_ and *α*_*j*_ respectively). Thus, we can model the contact frequency between bins *i* and *j* as 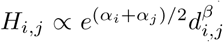 Finally, it is shown that the contact frequency between two bins *i* and *j* drops rapidly if there is a TAD boundary/insulator (e.g., a binding site of CTCF proteins) [4, 17] between these bins. We capture this drop off property in our model to achieve a more accurate contact frequency prediction. More specifically, we introduce a resistance variable for every bin *k* as *r*_*k*_, to model reduction of contacts between any two bins *i* and *j* where *i < k < j* indicates potential existence of a separator/insulator at the bin *k*. Note that these variables are set utilizing solely the input Hi-C matrix. Thus, the Hi-C contact map frequency model based on the (physical) distance *d*_*i,j*_ can be summarized as follows

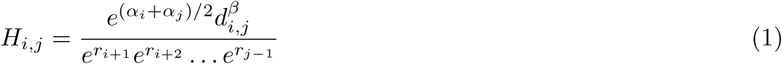

We denote this model for predicting Hi-C contact map frequencies as *distance based model*.

##### 5.1.2 Parameter optimization

The distance based model represented by equation 1 provides a framework for predicting the contact map frequencies after the deletion for any pair of bins *i* and *j* given the physical distance *d*_*i,j*_ between them. However, for this model to work accurately we need to first calculate the parameters *α*_*i*_, *r*_*i*_ for every bin *i* and the power-law coefficient *β* for the genome. We will estimate the parameter values of the above model to make the observed contact frequencies *H*_*i,j*_ and what is calculated based on the model are as close as possible. However, as the contact frequencies between different bins can be orders of magnitude different, instead of minimizing the absolute differences we try to make their ratio as close to 1 as possible (i.e. 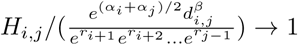). We can achieve this by minimizing the absolute log value of the above ratio as shown below

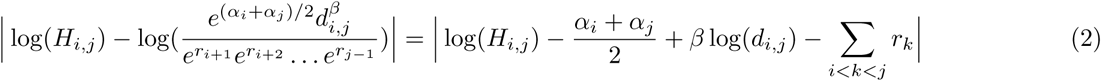

Note that *d*_*i,j*_ needs to satisfy the triangle inequality. Thus, we can estimate all parameters to model the contact frequency matrix *H* by solving the optimization problem below

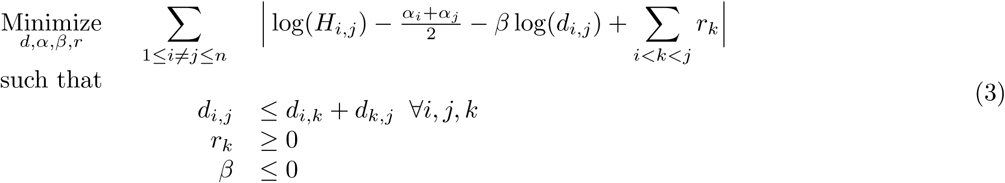

**Solving the optimization** The optimization problem in equation 3 is non-linear (because of the term *β* log(*d*_*i,j*_) and the triangle inequality can not be expressed in the log form). Instead of trying to solve the non-linear optimization problem, we are proposing a simplification for the optimization problem by pre-calculating some of the variables based on previous established models. We will first calculate the distance *d*_*i,j*_ using the chromosomal contacts and replace the calculated values in equation 3 to discover the optimal parameters (i.e., *α*_*i*_, *r*_*i*_ for every bin *i* and *β*). The 3-dimensional genome reconstruction and distance calculation using chromosomal contacts from Hi-C data is an active area of research with very promising results. Recently a model based on the shortest-path was introduced for reconstructing the 3-dimensional genome [45]. We utilize a similar idea to calculate the distances *d*_*i,j*_ for every pair of *i* and *j* given the Hi-C contact frequency matrix. First, we construct a graph where each node is a bin and we assume an edge between every pair of two nodes (corresponding to two bins *i* and *j*) and this edge is assigned a weight 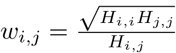 [45]. Second, the distance *d i,j* is calculated as the length of the shortest path between two nodes *i* and *j* on the constructed graph [45]. Note that this estimation guarantees that *d*_*i,j*_ always satisfies the triangle inequality as needed in our optimization formulation 3. We then fix the values of *d*_*i,j*_ calculated using this shortest-path strategy in the equation 3. Finally, by introducing slack variables *z*_*ij*_ we can turn the optimization problem 3 into a linear programming problem (LP) as shown below:

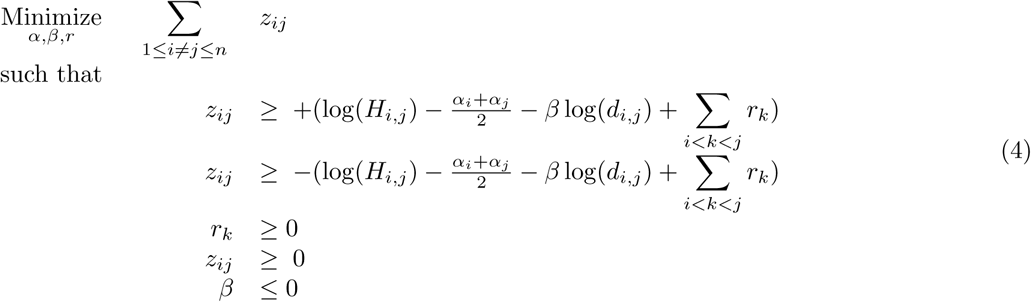

Note that in equation 4, *d*_*i,j*_ are pre-calculated using the shortest-path strategy and they become constants in the LP.

##### 5.1.3 Prediction of the contact frequencies

The parameters *α*_*i*_, *r*_*i*_ (for every bin *i*) and *β* are calculated using the LP optimization (4) and we assume they will not change as a result of the deletion. However, the distance *d*_*i,j*_ between bins *i* and *j* can be changed significantly due to the deletion Δ_*x,y*_ if *i < × < y < j*. Thus, we approximate the new distance *d*_*i,j*_ between bins *i* and *j* after the deletion by calculating an upper bound of that distance. We are assuming the distance between two bins which are in the same side (upstream or downstream) of the deletion will not be changed as a result of the deletion. Note that after the deletion Δ_*x,y*_ the bins *x* − 1 and *y* + 1 will be adjacent and considering the triangle inequality of the distance metric *d*, we will calculate the upper bound of the distance *d*_*i,j*_ as follow:

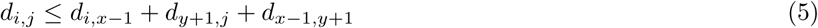

In the above formula, values *d*_*i,x*−1_ and *d*_*y*+1,*j*_ are previously calculated using the shortest-path strategy explained in the previous subsection and it does not change as a result of the deletion Δ_*x,y*_. Note that, in the new genome, bins *x* − 1 and *y* + 1 are now adjacent in the genome as such the distance *d*_*x*−1,*y*+1_ is negligible in comparison to other two terms in equation 5. We further use the average of the distances calculated for adjacent bins to approximate *d*_*x*−1,*y*+1_ (we also tested the prediction values assuming that *d*_*x*−1,*y*+1_ = 0 and observed almost identical results, mainly due to the fact that *d*_*x*−1,*y*+1_ was negligible in comparison to the other terms in equation 5). Finally, we use the upper bound distances calculated using equation 5 as the input to equation 1 to calculate the contact frequencies after deletion and use these values to fill the predicted contact frequencies *G*_*i,j*_ resulting from the deletion. In practice, we have shown that this approach is by far superior to using the genomic length based model to predict the contact frequency values (see figure S1).

#### 5.2 Estimating TAD fusion level of a deletion mutation

We denote the TAD fusion score of a deletion Δ_*x,y*_ as 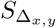 and define it as the difference between the expected number of genomic interactions between bins before and after deletion.

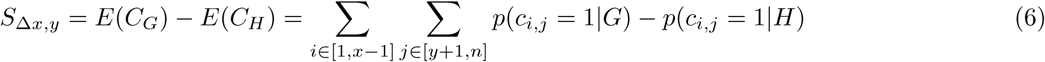

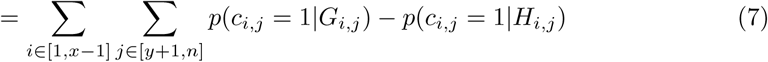

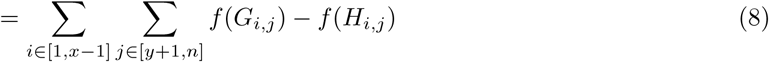

where random variable *C*_*G*_ and *C*_*H*_ represents the number of interactions between different bins assuming the Hi-C contact maps for the two genomes *H* and *G* respectively. The random variable *c*_*i,j*_ is an indicator variable representing existence of the interaction between bins *i* and *j* (i.e., *c*_*i,j*_ = 1 indicates an existence of interaction between bins *i* and *j* and *c*_*i,j*_ = 0 indicates the lack of such interaction in 3-dimensional space). We assume that the genomic interaction probability of a pair of bins only depends on their contact frequency. Therefore, we can use a function *f* to convert the contact frequency to the genomic interaction probability for both *G* and *H*. Here, we approximate *f* (*x*) as the fraction between the number of genomic interacting bin pairs that have a contact frequency *x* and the total number of all bin pairs that have contact frequency *x*. We use the HiCCUP loop set [31] to determine if a pair of bins has a genomic interaction. We assume that any two bins *i* and *j* that are inside the same loop will have a genomic interaction and if they belong to two separate loops then they are not interacting. Finally, for any *x* where *f* (*x*) *< f* (*x* − 1) we will modify *f* (*x*) to be equal to *f* (*x* − 1) to ensure that function *f* is a monotonically increasing function (see figure S3)

Notice that in our model in theory the deletion should increase the interaction between the remaining bins (i.e. *G*_*i,j*_ *≥ H*_*i,j*_), thus that score should be positive for any deletion. However, in our prediction we estimated the new values assuming the lower bound and the fact that data has the noise resulted from experimental measurement, thus the predicted contact frequency can in practice be smaller than the original contact frequency (i.e. before deletion). Hence, the calculated score by our method can be negative. We could simply impose a heuristic/ad-hoc rule to make sure *G*_*i,j*_ *≥ H*_*i,j*_, however as our goal is to rank SVs by their score, we decided to keep those negative scores as this is are more representative of how our method works.

#### 5.3 Implementation

The linear programming problem (equation 4) has *O*(*n*^2^) variables where *n* is the number of bins of a chromosome. Assuming for one human chromosome we have over 50, 000 bins (e.g. chromosome 1 with bin length 5kbp), this results in over 2 *×* 10^9^ variables. This large size problem is not efficiently solvable with our current LP solvers (e.g. CPLEX). Thus, we employed a strategy to speed up the running time of the method in practice by making some compromises as follow: (i) in regard to estimating the parameters, we partitioned each chromosome into smaller overlapping segments (6 Mbp segments with 3 Mbp overlaps) and estimated the parameters for each segment, (ii) in the objective function of the linear programming (equation 4) we only considered the summation of bin pairs that the genomic distance was at most 500 kbp apart (iii) Similarly, for calculating the shortest path between any two bins, we only considered bins that were inside the segment being considered (iv) The parameter *β* is limited to be in the range of the reported values in the literature (−2 *≤ β ≤* −1) [44, 49]. We have performed extensive simulation to illustrate the robustness of our method under different ranges of input parameters. Note that although larger values of these parameters provide a slightly more accurate result, these parameter settings make the running time of the method increase significantly. Thus, we are compromising the accuracy and efficiency in our parameter value selection.

## SUPPLEMENTARY MATERIAL

### 1.1 Evaluating the Hi-C contact frequency prediction

At the core of the proposed method to estimate the TAD fusion score is predicting the changes in contact frequencies as a result of the deletion. Thus, we first evaluate the accuracy of the proposed approach in section 5.1 for predicting contact frequencies, denoted as *distance based model* (Figure S1). Since we do not have any experimental Hi-C dataset of genomes with deletions, we evaluate the prediction power of this approach by predicting the interaction between two adjacent regions by only using the information about the interactions inside each region. More specifically, for a window size *w* and a bin *u*, we use all the contact frequencies *H*_*i,j*_ of the left region of the bin *u* (i.e. (*i, j*) *∈* [*u* − *w, u*]^2^) and the right region of the bin *u* (i.e. (*i, j*) *∈* [*u* + 1, *u* + *w* + 1]^2^) to predict the contact frequency values *H*_*i,j*_ where (*i, j*) *∈* [*u* − *w, u*] *×* [*u* + 1, *u* + *w* + 1] and compare with the real values *H*_*i,j*_ to evaluate the prediction accuracy. This evaluation is necessary for the later step where we estimate the TAD fusion score of a deletion since we also assume the deletion does not change the interaction inside the left region and the right region and we use that intra-region interaction to predict the inter-region interaction change due to the deletion. We compared our distance based model prediction accuracy (see section 5.1) with the one from the genomic length based model *H*_*i,j*_ *∝ e*^*α*^ *|i* −*j |*^*β*^. Note that, the genomic length based model is widely used in the literature [44, 49]. This comparison was done on full chromosome 21 (one of the shortest chromosomes) by sampling bins that are location in multiples of 25kbp. The result shows that our distance based model significantly outperforms the length based model (figure S1 and supplementary table **??**). For larger window sizes (*w* between 500kbp 2Mbp), this result is somewhat expected since the genomic length based model does not take into account the TAD structure while our distance based model does (parameters *r*_*i*_ in equation 4). However, for smaller window sizes (e.g. 125kb or 250kb), although the length based model can predict very well (average PCC = 0.82), our prediction using distance based model is still superior (average PCC = 0.91, see an illustrative example in Supplementary Figure S1). This result shows that our distance based model can not only predict the long range interaction by taking into account the TAD structure, but also can predict detail short range interaction better by using 3D structure features.

Furthermore, we also compared our distance based model for predicting contact frequencies between genomic pairs crossing several validated somatic deletions in K562 sample [50] and compared against the experimentally Hi-C matrix of the same sample [31]. The results of K562 comparison can be seen in Supplementary Figure S2 and Supplementary Table S2. In both these experiments we showed that our proposed *distance based model* outperforms the *genomic length based model*.

**Figure S1:**
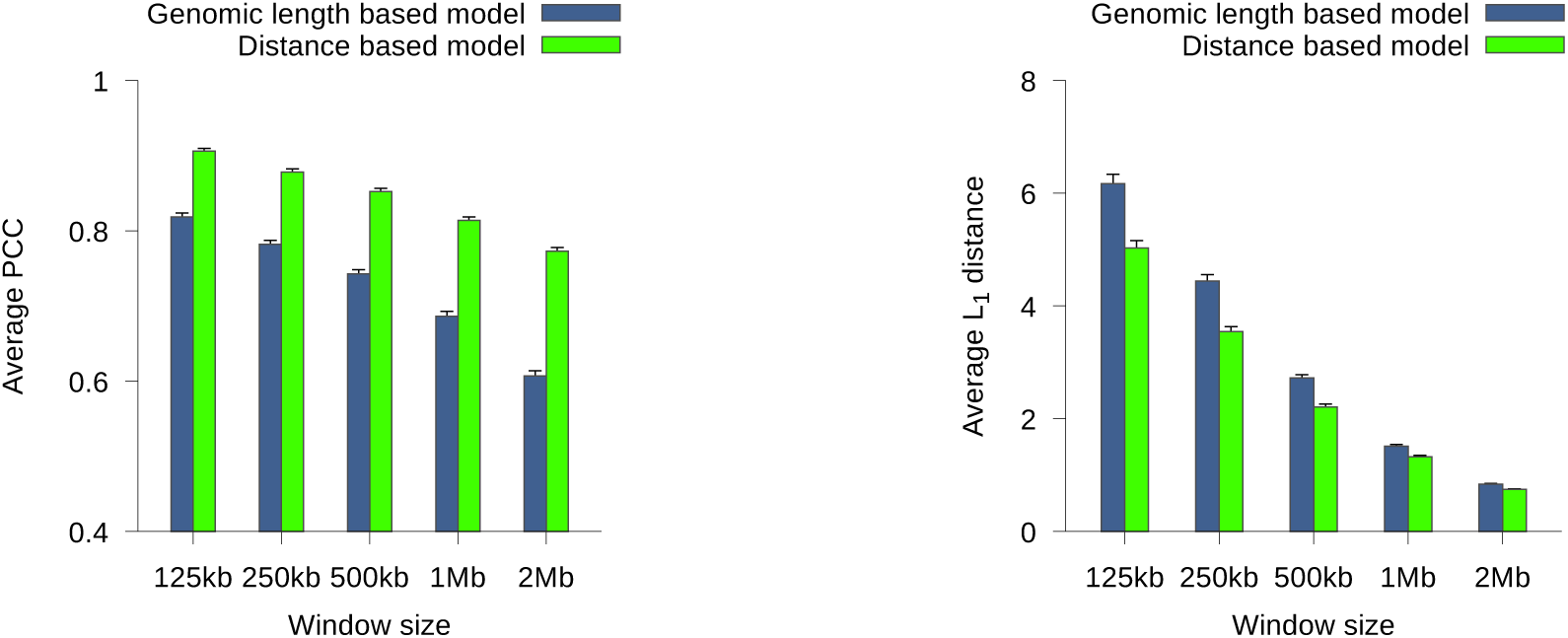
A comparison between the genomic length based model and the distance based model in predicting interregion interaction from intra-region interaction (i.e. upstream region and downstream region of a loci, we tried different region length values that we called window size, see section 5.1.1). We use two metrics including the Pearson correlation coefficient (PCC, left) and the *L*_1_ distance (right). For each metric, the value is the average of all values, taken at all bins that the location is a multiple of 25kbp on chromosome 21 (section 1.1). The error bar represents the standard error.

**Figure S2:**
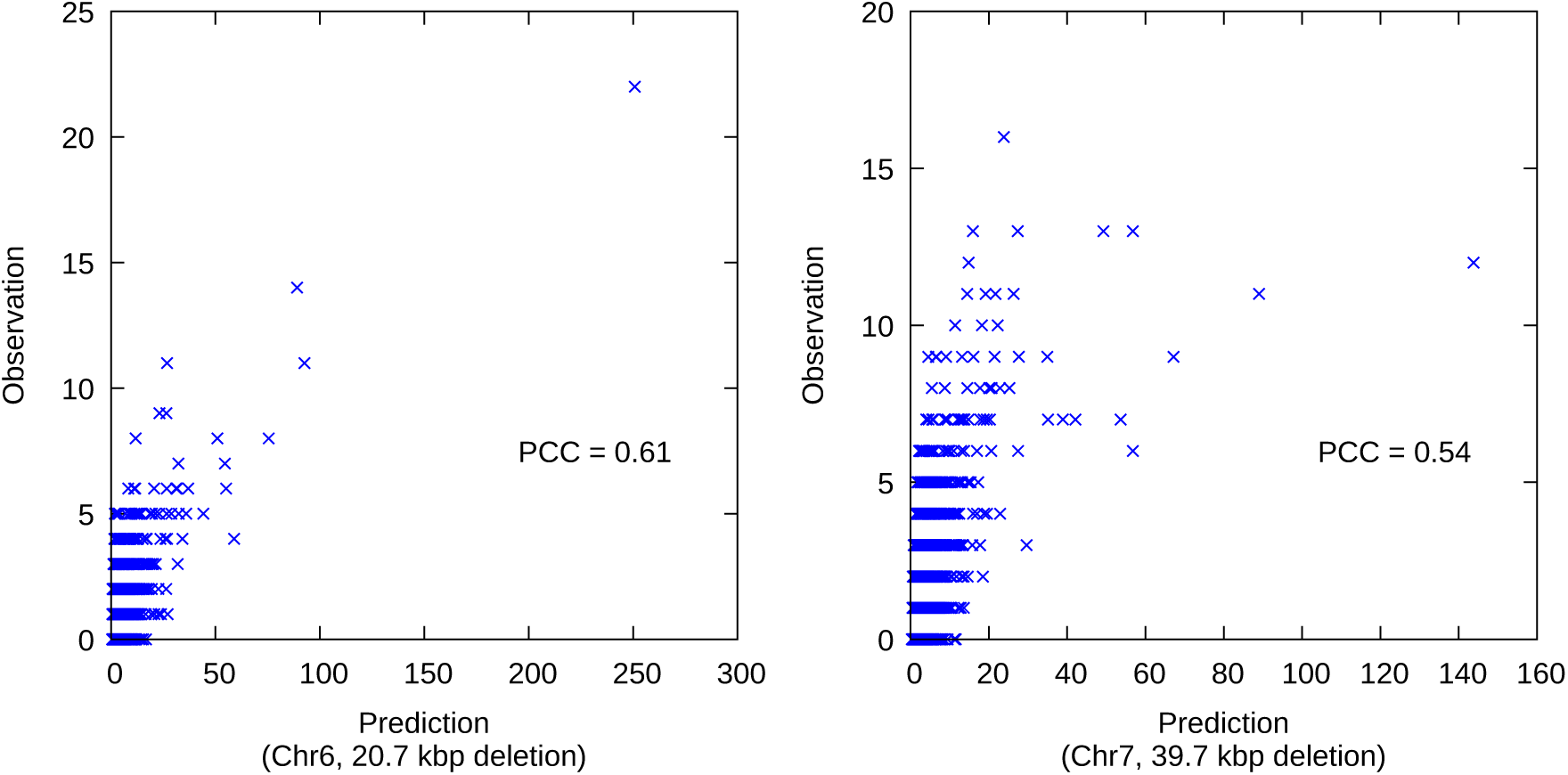
An illustration on the prediction of K562 Hi-C data around two longest deletions from K562 genome utilizing the GM12878 Hi-C data. Each data point represents the predicted value (x-axis) and the observed value (y-axis) of an interaction between a left loci (no longer than 250kbp in the upstream of the deletion) and a right loci (no longer than 250kbp in the downstream of the deletion).

**Figure S3:**
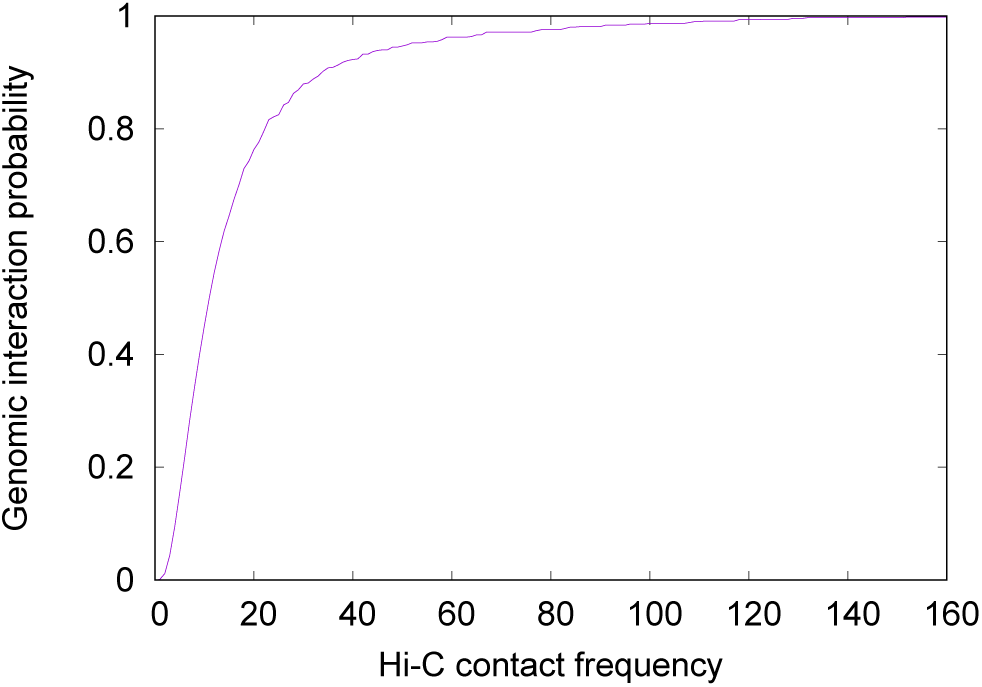
Genomic interaction probability as a monotonically increasing function of Hi-C contact frequency. This function is inferred from HiCCUP loops (of chromosome 1) of Rao et al [31].

**Figure S4:**
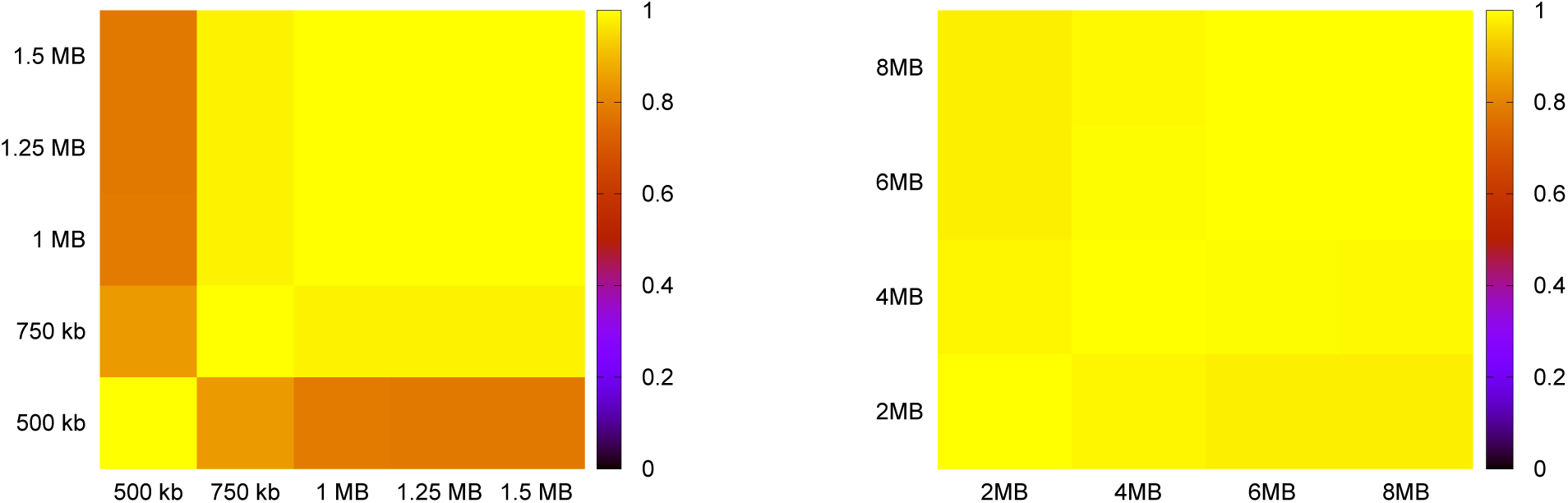
Evaluation of the robustness of our method when the parameter values are changed: Maximum interaction length (left) and segmentation length (right). The robustness is evaluated as the Pearson correlation coefficient between two vectors of the TAD-fusion scores of all 1KG deletions of chromosome 21 calculated with two different parameter values respectively.

**Table S1:**
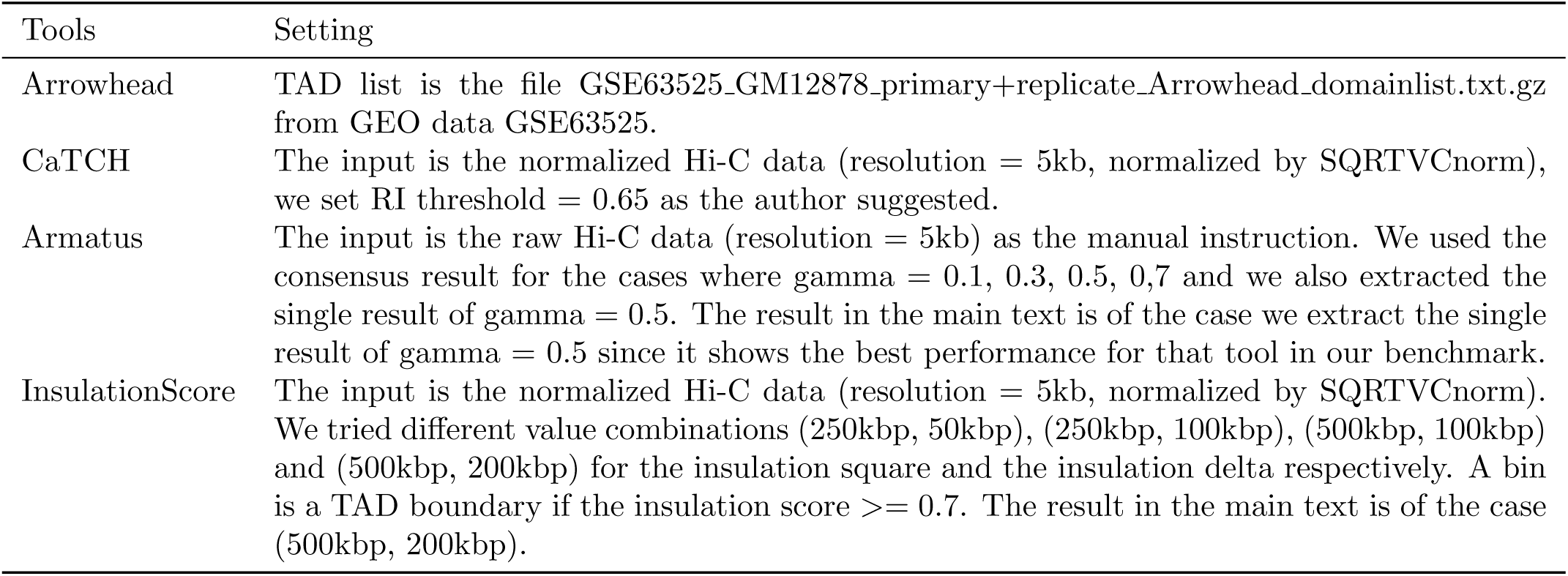
Parameter setting for TAD callers in the benchmark.

**Table S2:** A comparison on the prediction accuracy of K562 Hi-C data (as the Pearson correlation coefficient or *L*_1_ distance between predicted values and observed values) between the length based model and the distance based model.

| Window size | K562 deletion | | | | Pearson correlation coefficient | | $L_1$ distance | |
| --- | --- | --- | --- | --- | --- | --- | --- | --- |
|  | Chromosome | Left | Right | Length (bp) | Length based | Distance based | Length based | Distance based |
| 250 kbp | 4 | 10392430 | 10402181 | 9751 | 0.27 | 0.55 | 12 | 6 |
|  | 6 | 55825957 | 55846712 | 20755 | 0.52 | 0.61 | 5.7 | 3.4 |
|  | 7 | 8826500 | 8866251 | 39751 | 0.47 | 0.54 | 4.7 | 2.4 |
|  | 10 | 82879416 | 82891376 | 11960 | 0.43 | 0.48 | 6.9 | 6.1 |
|  | 11 | 48367528 | 48373693 | 6165 | 0.44 | 0.49 | 4.3 | 2.2 |
| 500 kbp | 4 | 10392430 | 10402181 | 9751 | 0.41 | 0.66 | 5.6 | 2.6 |
|  | 6 | 55825957 | 55846712 | 20755 | 0.41 | 0.48 | 3.6 | 1.7 |
|  | 7 | 8826500 | 8866251 | 39751 | 0.42 | 0.51 | 2.6 | 1.5 |
|  | 10 | 82879416 | 82891376 | 11960 | 0.37 | 0.41 | 5.3 | 2.8 |
|  | 11 | 48367528 | 48373693 | 6165 | 0.41 | 0.49 | 3 | 0.7 |
| 1 Mbp | 4 | 10392430 | 10402181 | 9751 | 0.56 | 0.67 | 3.8 | 1 |
|  | 6 | 55825957 | 55846712 | 20755 | 0.34 | 0.44 | 1.5 | 0.8 |
|  | 7 | 8826500 | 8866251 | 39751 | 0.36 | 0.56 | 1.2 | 0.6 |
|  | 10 | 82879416 | 82891376 | 11960 | 0.34 | 0.41 | 1.9 | 1.1 |
|  | 11 | 48367528 | 48373693 | 6165 | 0.39 | 0.47 | 1.7 | 0.3 |
| 2 Mbp | 4 | 10392430 | 10402181 | 9751 | 0.57 | 0.75 | 1.3 | 0.3 |
|  | 6 | 55825957 | 55846712 | 20755 | 0.32 | 0.36 | 0.8 | 0.3 |
|  | 7 | 8826500 | 8866251 | 39751 | 0.4 | 0.49 | 1.1 | 0.3 |
|  | 10 | 82879416 | 82891376 | 11960 | 0.33 | 0.35 | 0.9 | 0.3 |
|  | 11 | 48367528 | 48373693 | 6165 | 0.33 | 0.36 | 1.1 | 0.1 |

**Table S3:** A comparison on the number of TAD fusion deletions between the 1KG data and the permuted data for different top cut-off values. For each top cut-off value *n*, we extract the cut-off score as the score of the *n*^*th*^ deletion (sorted by the TAD-fusion score), then any permutation deletion that the score is greater than that cut-off score will be considered as a TAD-fusion deletion.

| Method |  | Observed number of<br>TAD-fusion deletions<br>(1KG data) | Expected number of<br>TAD-fusion deletions<br>(permuted data) | Z-score |
| --- | --- | --- | --- | --- |
| TAD-fusion score |  |  |  |  |
| cut-off = top 80 | | 80 | $257.6 \pm 14.2$ | -12.5 |
| cut-off = top 90 | | 90 | $269.9 \pm 14.5$ | -12.4 |
| cut-off = top 100 | | 100 | $296.9 \pm 15.1$ | -13.1 |
| cut-off = top 110 | | 110 | $321.7 \pm 15.6$ | -13.6 |
| cut-off = top 120 | | 120 | $332.4 \pm 15.9$ | -13.4 |

